# Alpha-synuclein-induced mitochondrial dysfunction is mediated via a sirtuin 3-dependent pathway

**DOI:** 10.1101/357624

**Authors:** Jae-Hyeon Park, Marion Delenclos, Ayman H. Faroqi, Natasha N. DeMeo, Pamela J. McLean

**Affiliations:** Department of Neuroscience, Mayo Clinic, Jacksonville, FL, USA; Mayo Clinic Graduate School of Biomedical Sciences, Mayo Clinic College of Medicine, Jacksonville, FL, USA

**Keywords:** α-Synuclein, Sirtuin 3, Mitochondria dysfunction, Parkinson’s disease

## Abstract

The sirtuins are highly conserved nicotinamide adenine dinucleotide (NAD^+^)-dependent enzymes that play a broad role in cellular metabolism and aging. Mitochondrial sirtuin 3 (SIRT3) is downregulated in aging and age-associated diseases such as cancer and neuro-degeneration and plays a major role in maintaining mitochondrial function and preventing oxidative stress. Mitochondria dysfunction is central to the pathogenesis of Parkinson disease with mutations in mitochondrial-associated proteins such as PINK1 and parkin causing familial Parkinson disease. Here, we demonstrate that the presence of alpha-synuclein (αsyn) oligomers in mitochondria induce a corresponding decrease in mitochondrial SIRT3 activity and decreased mitochondrial biogenesis. We show that SIRT3 downregulation in the presence of αsyn accumulation is accompanied by increased phosphorylation of AMP-activated protein kinase (AMPK) and cAMP-response element binding protein (CREB), as well as increased phosphorylation of dynamin-related protein 1 (DRP1) and decreased levels of optic atrophy 1 (OPA1), which is indicative of impaired mitochondrial dynamics. Treatment with the AMPK agonist 5-aminoimidazole-4-carboxamide-1-β-d-ribofuranoside (AICAR) restores SIRT3 expression and activity and improves mitochondrial function by decreasing αsyn oligomer formation. The accumulation of αsyn oligomers in mitochondria corresponds with SIRT3 down-regulation not only in an experimental cellular model, but also *in vivo* in a rodent model of Parkinson disease, and importantly, in human post mortem brains with neuropathologically confirmed Lewy body disease (LBD). Taken together our findings suggest that pharmacologically increasing SIRT3 levels will counteract αsyn-induced mitochondrial dysfunction by normalizing mitochondrial bioenergetics. These data support a protective role for SIRT3 in Parkinson disease-associated pathways and reveals significant mechanistic insight into the interplay of SIRT3 and αsyn.

## Abbreviations

SIRT3: Sirtuin 3
α-synuclein: αsyn
AMPK: adenosine monophosphate activated protein kinase
CREB: cAMP-response element binding protein
DRP1: dynamin-related protein 1
OPA1: optic atrophy 1
HO-1: heme oxygenase-1
SOD2: superoxide dismutase 2
mtROS: mitochondrial reactive oxygen species.

## Introduction

Alpha-synuclein (αsyn) accumulation is believed to be a key step in the pathogenesis of Parkinson’s disease and related alpha-synucleinopathies. Despite predominant localization in the cytosol, αsyn is found localized to mitochondria in post-mortem Parkinson’s disease brain (Devi *et al.,* 2008). Mitochondrial accumulation of αsyn has been associated with impairment of complex-I dependent respiration, decreased mitochondria membrane potential, and increased levels of mitochondrial reactive oxygen species (mtROS) in multiple cellular models (Hsu *et al.,* 2000; Devi *et al.,* 2008; Reeve *et al.,* 2015; Ludtmann *et al.,* 2018). The evidence supporting a contribution of abnormal accumulation of αsyn to disruption of mitochondrial processes is compelling and indicates a crucial role for αsyn-induced mitochondrial dysfunction in Parkinson’s disease pathogenesis and alpha-synucleopathies.

The sirtuins (SIRT) are a family of nicotinamide adenine dinucleotide (NAD^+^)-dependent deacetylases and/or adenosine diphosphate (ADP)-ribosyltransferases that have long been recognized as essential for cell survival, metabolism, and longevity (Kyrylenko *et al.,* 2010). In mammals there are seven human SIRT homologs (SIRT1–7) with varied enzymatic activities. SIRT1, SIRT6, and SIRT7 predominantly reside in the nucleus whereas SIRT2 is located in the cytoplasm, and SIRT3, 4, and 5 reside in the mitochondria. SIRT3 is the predominant mitochondrial sirtuin and the major regulator of mitochondrial protein acetylation (Hebert *et al.,* 2013; Herskovits and Guarente, 2013; Gleave *et al.,* 2017). SIRT3 is expressed at high levels in the brain (Lombard *et al.,* 2007; Lopez-Otin *et al.,* 2013) and plays an important role in maintaining mitochondrial integrity, energy metabolism, and regulating mitochondrial oxidative pathways (Kong *et al.,* 2010; Bause and Haigis 2013). SIRT3 ‐mediated deacetylation activates enzymes responsible for the reduction of ROS production, such as superoxide dismutase 2 (SOD2) (Ansari *et al.,* 2017). Interestingly, SIRT3 acts as a pro-survival factor in neurons exposed to excitotoxic injury (Kim *et al.,* 2011) and recent studies demonstrate a neuroprotective effect of SIRT3 in cell culture models of stroke, Huntington’s disease, and Alzheimer’s disease (Fu *et al.,* 2012; Weir *et al.,* 2012; Yin *et al.,* 2015). Importantly, and relevant to the present study, overexpression of SIRT3 was recently demonstrated to prevent dopaminergic cell loss in a rodent model of Parkinson disease (Gleave *et al.,* 2017).

Experimental evidence supports SIRT3-induced protection against oxidative stress by enhancement of mitochondrial biogenesis and integrity (Liu *et al.,* 2017). The multifaceted mitochondrial health-enhancing capabilities of SIRT3 thus make it an attractive therapeutic target for neurodegenerative diseases where mitochondrial dysfunction is believed to contribute to disease pathogenesis. Herein, we investigate a role for SIRT3 in Parkinson disease progression and identify a potential mechanistic interaction between SIRT3 and oligomeric forms of αsyn. We hypothesize that mitochondrial αsyn reduces SIRT3 deacetylase activity and contributes to mitochondrial dysfunction and pathogenesis in Parkinson disease and related αsynucleinopathies. The data presented herein significantly advances our mechanistic understanding of SIRT3 in mitochondrial dysfunction and validates a protective role for SIRT3 in Parkinson disease. Overall we confirm the potential application of SIRT3 activators as prospective targets for pharmacological strategies against neurodegeneration in Parkinson disease and related alpha-synucleinopathies.

## Materials and methods

### Cell culture

A stable cell line co-expressing human αsyn fused to either the amino-terminal (SL1) or carboxy-terminal fragment (SL2) of humanized *Gaussia princeps* luciferase was generated and described previously (Moussaud *et al.,* 2015). H4 SL1&SL2 cells were maintained at 37°C in a 95% air/5% CO2 humidified incubator in Opti-MEM supplemented with 10% FBS. Stock cultures were kept in the presence of 1μg/ml tetracycline (Invitrogen) to block the expression of the transgenes (SL1&SL2). αSyn expression is turned on or off by the absence (Tet- cells) or presence (Tet+ cells) of tetracycline respectively.

### Rodent Stereotaxic surgery

Adult female Sprague Dawley rats (225-250g, Envigo, USA) were housed and treated in accordance with the NIH Guide for Care and Use of Laboratory animals. All animal procedures were approved by the Mayo Institutional Animal Care and Use Committee and are in accordance with the NIH Guide for Care and Use of Laboratory animals. All viral vector delivery surgical procedures and tissue processing was performed as previously described by our group (Delenclos *et al.,* 2016). Briefly, adeno-associated-virus (AAV) serotype2/8 expressing human αsyn fused with either the C-terminus (AAV-SL1) or N-terminus (AAV-SL2) of *Gaussia princeps* luciferase was produced by plasmid triple transfection with helper plasmids in HEK293T cells. 48 hours later, cells were harvested and lysed in the presence of 0.5% sodium deoxycholate and 50U/ml Benzonase (Sigma-Aldrich, St. Louis, MO) by freeze-thawing, and the virus was isolated using a discontinuous iodixanol gradient. The genomic titer of each virus was determined by quantitative PCR. A combination of AAV-SL1 (8.1012gc/ml) + AAV-SL2 (8.1012 gc/ml) was delivered directly to the right substantia nigra (SN) using stereotaxic surgery (coordinates: AP −5.2mm, ML +2.0mm, DV +7.2mm from dura) (Paxinos and Watson, 1998). AAVs were infused at a rate of 0.4μL/min (final volume 2μL) using a microinjector (Stoelting). A group of control animals were injected with 2μL of AAV8 expressing full length of humanized *Gaussia princeps* luciferase (AAV8-Hgluc).

### Human brain tissue

Frozen human post-mortem brain was provided by the Mayo Clinic brain bank at the Mayo Clinic in Jacksonville. For this study, striatum (STR) samples from 10 control patients (6 females, 4 males) and 10 patients diagnosed with Lewy body disease (4 females and 6 males) were included. Detailed information of brain tissues is provided in Table 1. Each frozen brain sample was weighed and homogenized in 10X volume of RIPA buffer (50mM Tris–HCl, pH 7.4, 150mM NaCl, 1mM EDTA, 1mM EGTA, 1.2% Triton X-100, 0.5% sodium deoxycholate, and 0.1% SDS) containing 1mM phenylmethylsulfonyl fluoride (PMSF), protease inhibitor cocktail, and halt phosphatase inhibitor cocktail, followed by sonication and centrifugation for 15 min at 16,000 × g at 4 °C to remove cellular debris. Supernatants were collected, protein concentration was determined by Bradford assay, and samples were processed for immunoblotting.

**Table 1.**
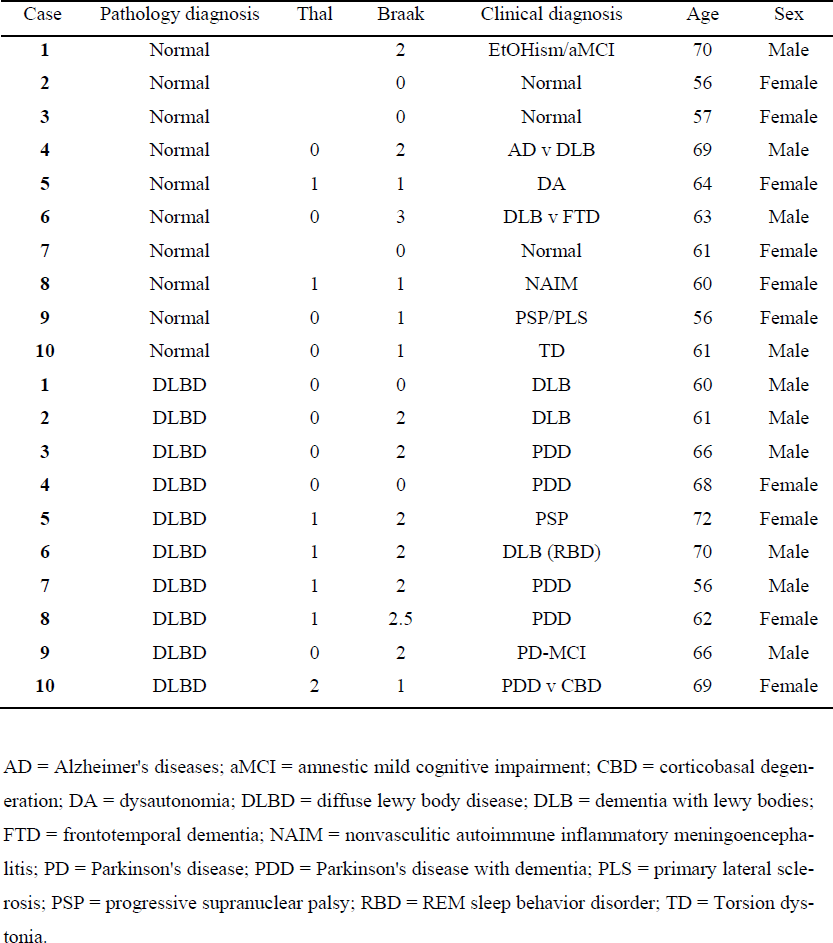
Human brain samples.

### Immunofluorescence

Cells were cultured on 12-mm glass coverslips with or without 1μg/ml tetracycline for 72h. Cells were washed with phosphate-buffered saline (PBS) and incubated with 300nM with MitoTracker-Green (Molecular Probes, Inc., Eugene, OR, USA) according to the manufacturer’s protocol to visualize mitochondria. Cells were fixed with 4% paraformaldehyde for 10min at room temperature (RT) and washed three times in 1X Tris-buffered saline (TBS) (500mM NaCl, 20 mM Tris, pH 7.4), blocked for 1h in 1.5% goat serum, 0.5% Triton X-100 in 1X TBS and incubated overnight at 4°C with primary antibodies (SIRT3 and human αsyn). The following day cells were washed and treated with Alexa Fluor ^®^ 488 and 568 secondary antibodies for 1h at RT (see Table 2, for details of the antibodies used in the study). Coverslips were mounted on Super Frost Plus slides with Vectashield Hardset (Vector Labs, Burlingame, CA) and cells were visualized using an Axio observer inverted microscope (Carl Zeiss, Germany).

**Table 2.** Antibodies used for western blot and immunocyhistochemistry.

| Antibody | Source | Dilution |
| --- | --- | --- |
| α-Synuclein (mouse) | BD Transduction Laboratories (61078) | 1:2000 (WB) |
|  | Biolegend (SIG39730) | 1:2000 (WB, ICC) |
| OPA1 (mouse) | BD Transduction Laboratories (612606) | 1:2000 (WB) |
| SIRT3 (mouse) | Santa Cruz (sc-135796) | 1:1000 (WB) |
| SIRT3 (rabbit) | Novus Biologicals (NBP1-31029) | 1:1000 (WB) |
|  |  | 1:500 (ICC) |
| Heme oxygenase-1 (rabbit) | Cell Signaling (5853s) | 1:1000 (WB) |
| AMPK (rabbit) | Cell Signaling (5831T) | 1:1000 (WB) |
| Phospho-AMPK (rabbit) | Cell Signaling (2535s) | 1:1000 (WB) |
| CREB (rabbit) | Cell Signaling (4820s) | 1:1000 (WB) |
| Phospho-CREB (rabbit) | EMD Millipore (06-519) | 1:2000 (WB) |
| DRP1 (rabbit) | Bethyl Laboratories (A303-410A-M) | 1:2000 (WB) |
| Phospho-DRP1 (rabbit) | Cell Signaling (3455s) | 1:1000 (WB) |
| SOD2 | abcam (ab13533) | 1:5000 (WB) |
| SOD2 (acetyl K68) (rabbit) | abcam (ab137037) | 1:2000 (WB) |
| COX IV (rabbit) | Cell Signaling (4850s) | 1:1000 (WB) |
| GAPDH (rabbit) | Santa Cruz (sc-25778) | 1:4000 (WB) |
|  | Abgent (AP7873a) | 1:4000 (WB) |
| Alexa Fluor 488 (goat anti-mouse) | Thermo Fisher Scientific (A11001) | 1:500 (ICC) |
| Alexa Fluor 568 (goat anti-rabbit) | Thermo Fisher Scientific (A11011) | 1:500 (ICC) |
| Goat anti-mouse HRP | Southern biotech (1010-05) | 1:5000 (WB) |
| Goat anti-rabbit HRP | Southern biotech (4010-05) | 1:5000 (WB) |

WB = western blot; ICC = immunocytochemistry.

### Gaussia luciferase protein-fragment complementation assays

Luciferase activity was measured in 15ug of cell lysate in a multilabel plate reader (EnVision, PerkinElmer; Waltham, MA, USA) following the injection of the cell permeable substrate, coelenterazine (20mM, NanoLight).

### Western blotting analysis

To prepare whole cell lysates, cells were washed twice with ice-cold PBS and total proteins were isolated by incubating the cells on ice in radio-immunoprecipitation assay (RIPA) lysis buffer (50mM Tris–HCl, pH 7.4, 150mM NaCl, 1mM EDTA, 1mM EGTA, 1.2% Triton X-100, 0.5% sodium deoxycholate, and 0.1% SDS) containing 1mM phenylmethylsulfonyl fluoride (PMSF), protease inhibitor cocktail, and halt phosphatase inhibitor cocktail. Collected cells were sonicated on ice and centrifuged at 10,000 × g for 10min at 4°C. The protein concentration was determined with Bradford reagent. 15μg proteins were separated on Bis-Tris polyacrylamide gradient gels (NuPAGE Novex 4-12% Bis-Tris Gel, Life tech) and transferred to nitrocellulose (NC) membranes. Membranes were then blocked for 1h at RT in TBST (500mM NaCl, 20mM Tris, 0.1% Tween 20, pH 7.4) supplemented with 10% non-fat dried milk. Subsequently membranes were incubated overnight at 4°C with primary antibodies followed by 1h at RT with HRP-conjugated secondary antibodies (Table 2). Proteins were detected using an enhanced chemiluminescent detection system (ECL, EMD Millipore) and a CCD imaging system (LAS-4000, Fujifilm, Japan).

### Mitochondria/cytosol fractionation

Cells were lysed in buffer A (0.25 M sucrose, 10mM Tris–HCl [pH 7.5], 10 mM KCl, 1.5 mM MgCl2, 1 mM EDTA, 1 mM dithiothreitol, and 0.1 mM PMSF) with an homogenizer. Homogenates were centrifuged at 700 × g for 5min at 4°C, and supernatants were collected and centrifuged at 10,000 × g for 30min at 4°C. The supernatants were used as the cytosolic fraction, and the pellet was used as the mitochondrial fraction. The pellets were resuspended in buffer B (0.25M sucrose, 10mM Tris–HCl [pH 7.5], 10mM KCl, 1.5mM MgCl_2_, 1mM EDTA, 1 mM dithiothreitol, 0.1mM PMSF, and 1% NP 40). To confirm the purity of the mitochondrial fraction, the lysates were probed for the specific mitochondria marker cytochrome *c* oxidase IV (COXIV).

### Isolation of rat brain mitochondria

Striatum (STR) and midbrain containing substantia nigra (SN) were dissected and homogenized in 0.5mL of ice-cold MIBA (10mM Tris–HCl [pH 7.4], 1mM EDTA, 0.2M D-mannitol, 0.05M sucrose, 0.5mM sodium orthovanadate, 1mM sodium fluoride and dissolved in water) containing 1X protease inhibitors and hand-held homogenizer for 40 strokes on ice. The homogenate was transferred into 1.5 mL tubes and then centrifuged at 500 × g for 5min. The pellet was discarded, and remaining supernatant was centrifuged at 11,000 × g for 20min at 4°C, yielding the heavy mitochondrial (HM, pellet) and the light mitochondrial (LM, supernatant) fraction. The HM pellet was washed twice with 1mL ice-cold MIBA buffer it was re-suspended in 0.1 - 0.3mL of MIBA to yield the final solution enriched in mitochondria.

### Mitochondrial respiration analysis

The oxygen consumption rate (OCR) was assessed using a Seahorse Bioscience XF96 analyzer (Seahorse Bioscience, Billerica, MA, USA) in combination with the Seahorse Bioscience XF Cell Mito Stress Test assay kit according to the manufacturer’s recommendations. H4 SL1&SL2 cells were seeded in 12-wells of a XF 96-well cell culture microplate (Seahorse Bioscience, 102601-100) and grown to 70% confluency in 200μL of growth medium prior to analysis. On the day of assay, culture media were changed to assay medium with 175μL (Dulbecco’s Modified Eagle’s Medium, D5030), supplemented with 25mM glucose, 2mM glutamine, and 2mM pyruvate. Prior to assay, plates were incubated at 37°C for 1h without CO_2_. Thereafter successive OCR measurements were performed consisting of basal OCR, followed by OCR level after the automated injection of 25μl oligomycin (20μM), 25μl carbonyl cyanide 4-(trifluoromethoxy) phenylhydrazone (FCCP) (20μM), and a combination of 25μl rotenone + antimycin A (12μM), respectively. After the assays, plates were saved and OCR was normalized to the total protein amount per well.

### SIRT3 siRNA transfection

Small interfering RNAs (siRNAs) for human SIRT3 (sc-61555, Santa Cruz Biotechnology, CA, USA) and control non-target siRNA (SN-1003, Negative Control, Bioneer, Daejeon, Korea) were reconstituted in siRNA buffer (Qiagen, CA) following the manufacturer’s instructions and transfections of SL1SL2 cells conducted using Lipofectamine 3000 reagent (Invitrogen, CA, USA). Briefly H4 SL1&SL2 cells were seeded in 6-well culture plate 24h before transfection. Subconfluent cells were treated either with SIRT3 siRNA (100nM) or non-targeting siRNA (20nM) complexed with Lipofectamine for 3h. The extent of knockdown was evaluated by western blot analysis.

### Determination of mitochondrial ROS

MitoSOX™ Red fluorescent probe (Molecular Probes, Inc., Eugene, OR, USA) was used to visualize mitochondrial superoxide production according to the manufacturer’s protocol. Briefly, H4 SL1&SL2 grown on 12-mm glass were washed twice with PBS to remove the medium and incubated with 2.5μM MitoSOX Red reagent in the dark at 37°C. Cells were washed gently three times with warm PBS buffer and imaged immediately after, under fluorescence microscopy.

### Statistical analysis

All data were analyzed by the Graph Pad Prism 7 software (San Diego, CA) and statistical significance was determined by one-way ANOVA analysis of variance with Tukey’s multiple comparisons test. Results presented as mean ± standard error of the mean (S.E.M.). For isolated mitochondria studies *in vivo*, a Mann-Whitney U test was used to analyze the Western blots, Differences were considered to be statistically significant with *P< 0.05, **P< 0.01, ^#^P< 0.05, ^##^P< 0.01, and n.s, not statistically significant (p > 0.05).

### Data Availability

Data sharing is not applicable to this article as no new data were created or analyzed in this study.

## Results

### Increased αsyn oligomers in mitochondria correlate with decreased SIRT3 protein levels

Although it has been described previously that αsyn localizes to mitochondria and αsyn over-expressing cells exhibit mitochondrial dysfunction (Devi *et al.,* 2008; Marongiu *et al.,* 2009; Nakamura *et al.,* 2011) the relationship between αsyn oligomers and mitochondria in pathologic conditions and the mechanisms whereby αsyn induces mitochondrial dysfunction are still poorly understood. Herein, we use a previously described inducible cell model of human αsyn overexpression that results in formation of intracellular oligomeric species over time (Moussaud *et al.,* 2015). This tetracycline-off (Tet-off) stable cell line facilitates monitoring of αsyn oligomerization *in situ* via a split luciferase protein–fragment complementation assay. To determine if αsyn oligomeric species are localized within mitochondria, cells were harvested at various time points after tetracycline removal, mitochondrial-enriched fractions were isolated, and luciferase activity was measured as a surrogate for αsyn oligomeric species. Luciferase activity increased in a time-dependent manner in both the mitochondrial (Fig. 1A and B) and cytosolic fractions (Suppl. Fig. 1A) of the cells. Increased αsyn oligomers were confirmed by the detection of increased high molecular weight species in both compartments 72h after removal of tetracycline (Suppl. Fig. 1B). GAPDH and COXIV immunoblotting confirmed purity of mitochondrial fractionation (Figs. 1A and B). Interestingly, the increase in mitochondrial-localized αsyn oligomers was accompanied by a decrease in SIRT3 protein levels beginning 12h after αsyn expression was turned on, and becoming significant by 24h (Fig. 1A). Immunocytochemistry confirmed decreased SIRT3 immunofluorescence in cells accumulating αsyn oligomers (Suppl. Fig. 1C, Tet– 72h) compared to control (Suppl. Fig. 1C, Tet+ 72h). In support of a αsyn-mediated effect on SIRT3 levels, knockdown of SIRT3 increased αsyn oligomers with a corresponding increase in αsyn protein levels (Figs. 1C and D).

**Figure 1:**
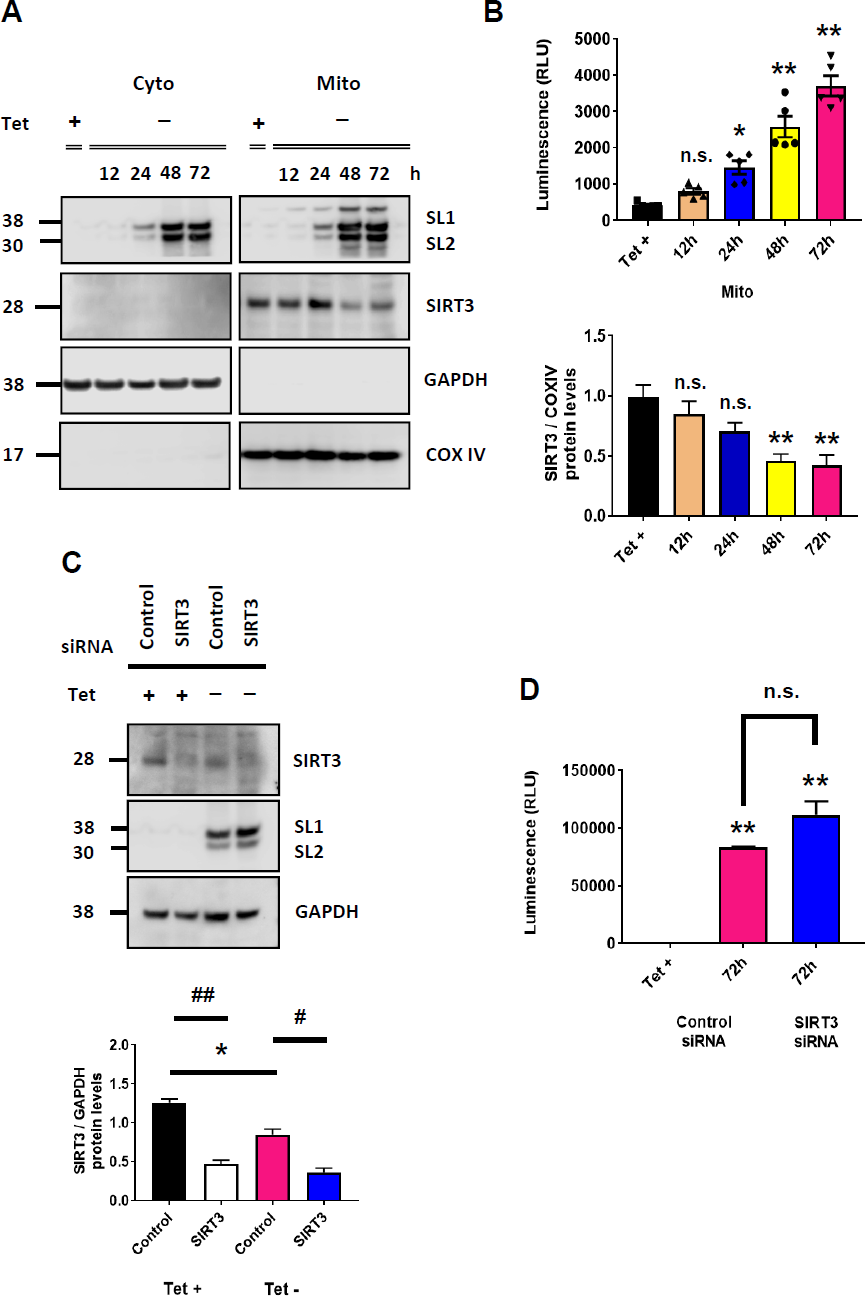
αSyn oligomers localize to mitochondria in H4 SL1&SL2 cells and induce a decrease in SIRT3 expression. (A) Representative cropped western blots showing αsyn and SIRT3 in cytosolic and mitochondrial fractions at different time points (0 – 72h) (B) Quantification of SIRT3 protein levels in mitochondria demonstrates significant decrease in SIRT3 after 24h. Increased αsyn oligomers at 24h and up to 72h are detected by luminescence assay (RLU: relative luciferase units), n=5. (C) H4 SL1&SL2 cells transfected with SIRT3 siRNA have less SIRT3 expression after 72h in whole cells lysates n=2 (D) Level of αsyn oligomers is significantly increased in cells transfected with SIRT3 siRNA. Error bars represent the mean ± S.E.M. *P < 0.05, **P < 0.01 compared to control conditions; #P < 0.05, ##P < 0.01 compared to SIRT3 siRNA transfection. n.s: not significant. In panel (A) αsyn and SIRT3 bands are obtained from different samples run on different gels. COXIV, GAPDH, and SIRT3 are all from same samples and blot. Loading controls for αsyn blot are not shown. In panel (C) αsyn, SIRT3, and GAPDH are probed on same blot.

### Mitochondrial oligomeric αsyn induces SIRT3 inactivation via AMPKα-CREB signaling pathway

SIRT3 regulates the synthesis of ATP by modulating AMP-activated protein kinase (AMPK), which acts as a sensor of cellular homeostasis. Cells with decreased SIRT3 function show reduced AMPKα phosphorylation (Shi *et al.,* 2005; Lombard *et al.,* 2007; Pillai *et al.,* 2010) and reduced phosphorylation and activity of cAMP response element binding protein (CREB). In addition, previous studies have shown that overexpression of αsyn reduces AMPKα activation in neuronal cells (Dulovic *et al.,* 2014). Because expression of αsyn oligomers in mitochondria results in decreased SIRT3 expression, we next examined the levels of p-AMPKα and p-CREB. At 72h, when SIRT3 expression is significantly decreased and mitochondrial αsyn oligomers are present (Fig. 1A), we detected a significant decrease in p-AMPKα (Thr172) and p-CREB (Ser133) (Fig. 2A and B). AMPKα-CREB signaling was also decreased in cells transfected with SIRT3 siRNA compared to control siRNA (Supplementary Fig. 2). To further validate modulation of the AMPKα-CREB signaling pathway by mitochondrial αsyn oligomers we asked whether treatment with 5-aminoimidazole-4-carboxamide-1-β-d-ribofuranoside (AICAR), an AMPKα agonist, could prevent αsyn-induced changes in mitochondrial SIRT3 and associated signaling proteins. SL1&SL2 cells were treated with 2 mM AICAR for 2h in accordance with a previous study (Takeuchi *et al.,* 2013), and harvested 72h after tetracycline removal. A significant increase in p-AMPKα and p-CREB levels was observed (Fig. 2A, #P<.0.05) and importantly, led to a partial restoration of SIRT3 levels to control levels (Fig. 2B, #P<0.05) and significantly decreased the level of αsyn oligomers (Fig. 2C). Because SIRT3 is a deacetylase known to modulate the acetylation of SOD2 (Qiu et al., 2010), we examined the level of acetylated SOD2 in cells overexpressing αsyn oligomers. Acetylated SOD2 (K68) was significantly increased in cells overexpressing αsyn compared to control (Fig. 2D, **P<0.01), consistent with reduced SIRT3 levels and activity. Of note, αsyn overexpression had no effect on total SOD2 levels which remained consistent in all conditions (Fig. 2D).

**Figure 2:**
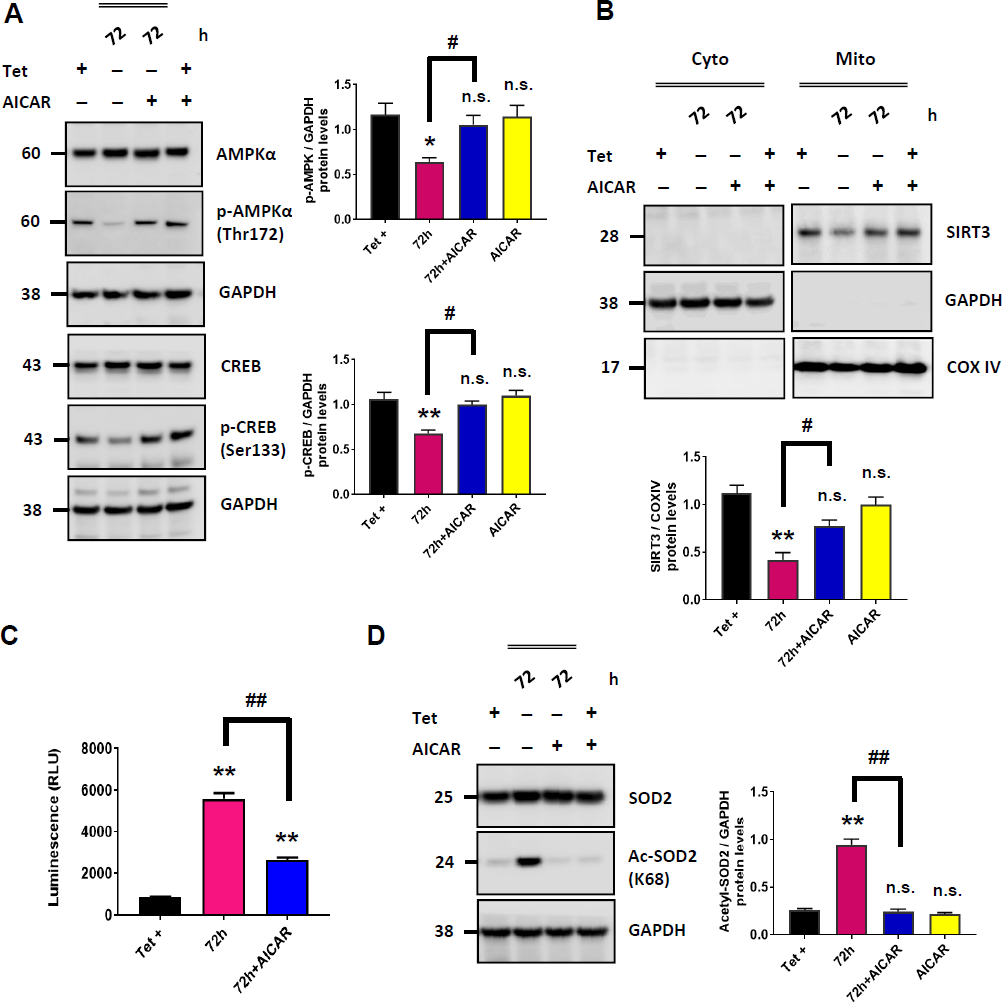
AICAR activates AMPK-CREB signaling pathway and increases SIRT3 activity to reduce αsyn oligomers. (A) Representative cropped western blots from showing AMPKα, p-AMPKα (Thr 172), CREB, and p-CREB (Ser 133) in H4 SL1&SL2 cells with/without 2mM AICAR. Quantification of blots show decreased of p-AMPKα (n=4) and p-CREB (n=3) at 72h, restored by AICAR treatment. (B) Levels of SIRT3 are restored after AICAR-treatment, n=3. (C) Luciferase assay shows activation of SIRT3 by AICAR significantly decreases αsyn oligomers (n=5). (D) Representative cropped western blot showing increased Ac-SOD2 (acetyl K68) with no change in total SOD2 in whole cells lysates. AICAR reversed the change in acetylated SOD2 level, n=3. Error bars represent the mean ± S.E.M (n = 3-5). *P < 0.05, **P < 0.01 compared to control conditions; #P < 0.05, ##P < 0.01 compared to AICAR treatment. n.s: not significant. In panel (A) the same samples were run on different gels and probed separately for AMPKα, p-AMPKα, and GAPDH, and CREB, p-CREB, and GAPDH respectively. In panels (B) and (D) separate blots were probed for SIRT3 and GAPDH, SIRT3 and COXIV, or SOD2, Ac-SOD2, and GAPDH.

### SIRT 3 activation attenuates αsyn-induced mitochondrial ROS

Because SIRT3 plays a crucial role in modulating ROS and limiting the oxidative damage of cellular components (Torrens-Mas *et al.,* 2017), we asked whether mitochondrial αsyn oligomers induce oxidative stress that can be rescued with SIRT3 activation. Cells overexpressing αsyn were stained with mitotracker red to visualize mitochondria and MitoSOX to monitor mitochondrial ROS production. Fluorescence microscopy revealed increased ROS at 72h compared to control condition (Tet+ 72h) (Fig. 3A). As predicted, AICAR-treatment reduced ROS production (Fig. 3A, bottom row). Increased oxidative stress and ROS can induce the expression of heme oxygenase-1 (HO-1) (Bansal *et al.,* 2013) and increased HO-1 mRNA and protein expression have been reported in a wide spectrum of diseases including neurodegenerative diseases such as Parkinson disease (Shipper *et al.,* 1998; Song *et al.,* 2009). In line with these data, we found a significant increase of HO-1 in cells expressing αsyn for 72h (Fig. 3B), and a concomitant decrease of HO-1 in cells treated with AICAR compared to control (Tet+ 72h) (Fig. 3B, P= n.s).

**Figure 3:**
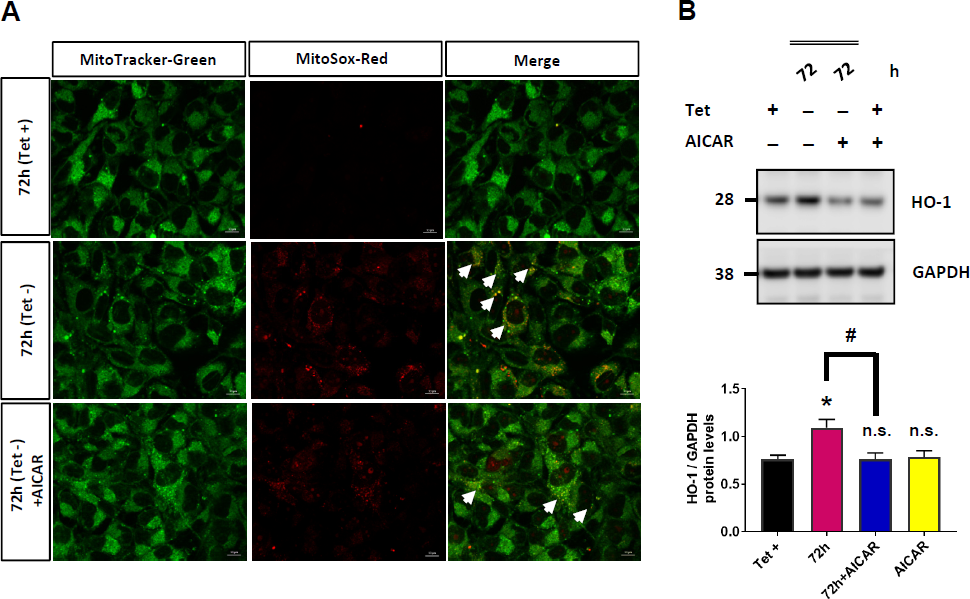
Activation of SIRT3 by AICAR attenuates ROS production. (A) Fluorescence microscopy images of MitoSox and Mitotracker staining in fixed H4 SL1&SL2 cells. αSyn oligomer expression increases mtROS at 72h, and AICAR treatment attenuates mtROS. Representative image from 3 experiments. MitoTracker-Green (mitochondria; green); MitoSox-Red (mitochondria; red); merged images (yellow). Scale bar = 10μm. White arrow indicates mtROS. (B) Representative cropped western blot of HO-1 and GAPDH in whole cells lysates from H4 SL1&SL2 cells. HO-1 level increases at 72h and is reduced after AICAR-treatment, n=5. Error bars represent the mean ± S.E.M. *P < 0.05 compared to control conditions; #P < 0.05 compared to AICAR treatment. n.s: not significant.

### αSyn impairs mitochondrial dynamics and bioenergetics which can be rescued by activation of SIRT3

Mitochondrial dynamics (fission/fusion) play a critical role in maintaining mitochondrial health, with the balance between fission (DRP1) and fusion (OPA1) proteins being crucial for neuronal function and survival. Changes in the expression and/or localization of fission/fusion proteins can impair this process and induce cell death. To determine the effect of αsyn oligomers on mitochondrial dynamics, we examined the expression of DRP1 and OPA1. In the presence of αsyn oligomers, DRP1 is recruited from the cytosol to the mitochondria (Fig. 4A). Consistent with the fact that phosphorylation of DRP1 at serine 616 activates mitochondrial fission, we detected increased levels of p-DRP1 in cells overexpressing αsyn (Suppl. Fig. 3). By contrast, OPA1 protein levels decreased over time in the mitochondrial fraction (Fig. 4A), consistent with a decrease in mitochondrial fusion. Several lines of evidence suggest that a decrease in OPA1 and the translocation of DRP1 to the mitochondria are crucial events that lead to mitochondrial fragmentation. When we evaluated the effect of AICAR on mitochondria dynamics we found OPA1 level in mitochondria restored (Fig. 4B) and phosphorylation of DRP1 significantly decreased (Fig. 4C). To determine if accumulation of αsyn in the mitochondria affects cellular bioenergetics we measured the oxygen consumption rate (OCR) in cell lysates using the Seahorse XF96 analyzer. The OCR was measured under basal conditions followed by the sequential addition of oligomycin (ATP synthase inhibitor), carbonyl cyanide 4-(trifluoromethoxy) phenylhydrazone (FCCP; mitochondrial uncoupler), and rotenone plus antimycin A (Complex I and III inhibitor) to assess ATP production, maximal respiration, and spare capacity respectively. Cells overexpressing αsyn had significantly decreased OCR in all paradigms tested when compared to control cells (Tet+) (Figs. 5A - E). This is highly suggestive of a mitochondria respiratory deficit in the presence of mitochondrial αsyn oligomers. We next analyzed the OCR of cells treated with AICAR and found that AICAR treatment was able to significantly restore the OCR level of basal respiration (Fig. 5B, #P <0.05) to the level of control and partially rescue ATP production and maximal respiration (Figs. 5C and D) with a trend toward restoration of spare capacity observed (Fig. 5E). Taken together, our data support a hypothesis whereby increased mitochondrial αsyn results in decreased mitochondrial function via a SIRT3-dependent cascade of events that can be rescued by restoring SIRT3 levels using an AMPKα agonist.

**Figure 4:**
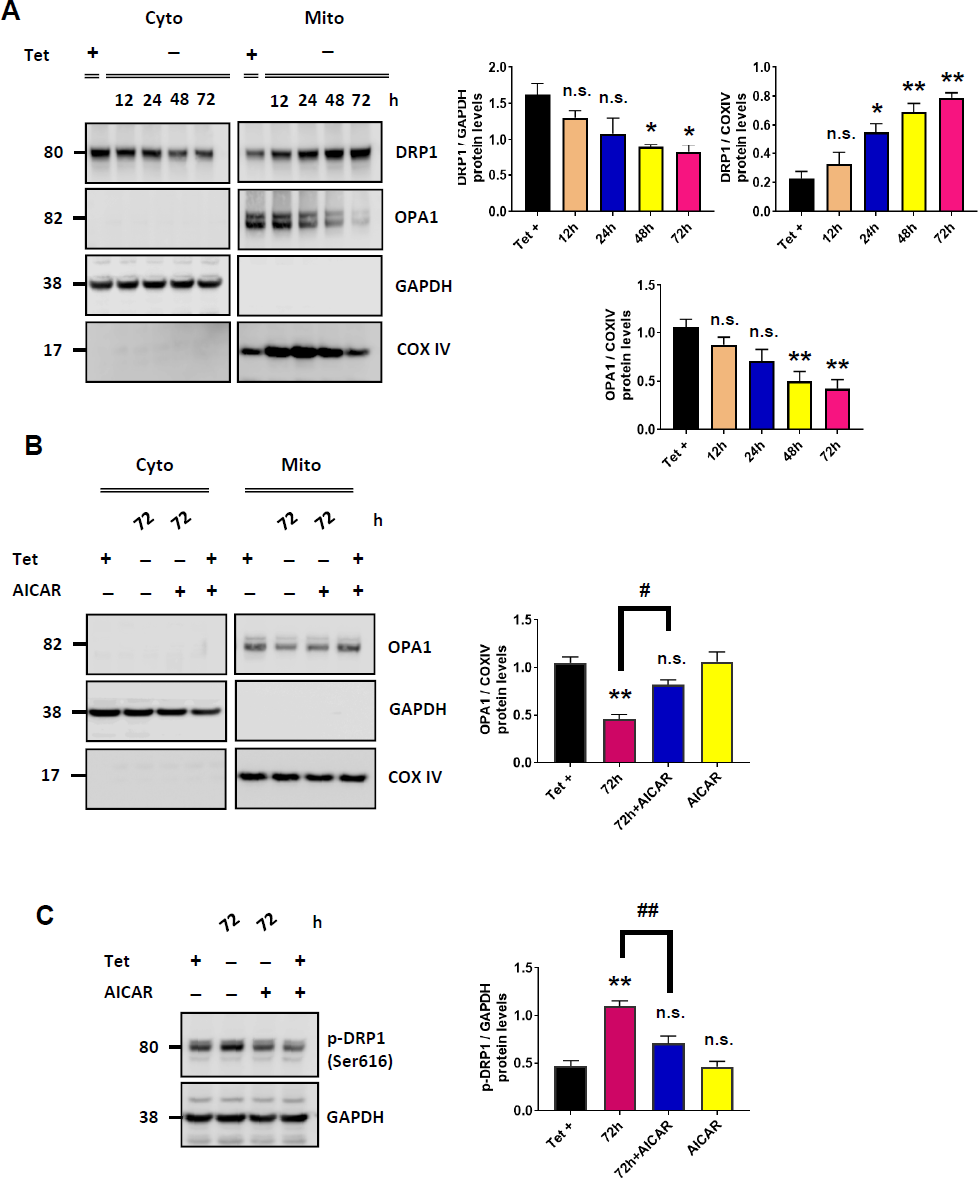
SIRT3 activation restores DRP1 and OPA1 expression and rescues impaired mitochondrial dynamics. (A) Representative cropped western blot showing DRP1 and OPA1 in cytosol and mitochondria from H4 SL1&SL2 cells over time (0 – 72h). (B) Quantitation of protein levels for DRP1 and OPA1 in cytosol and mitochondria from n=3 (DRP1) and n=4 (OPA1) blots. DRP1 and OPA1 bands were normalized to respective loading controls GAPDH and COXIV. (C and D) Representative cropped western blot from showing OPA1 (n= 3) and p-DRP1 (n=4) levels in cells treated with or without AICAR. AICAR-treatment restores the OPA1 levels in mitochondria and decreases p-DRP1 protein levels at 72h. Error bars represent the mean ± S.E.M. *P < 0.05, **P < 0.01 compared to control conditions; #P < 0.05, ##P < 0.01 compared to AICAR treatment. n.s: not significant.

**Figure 5:**
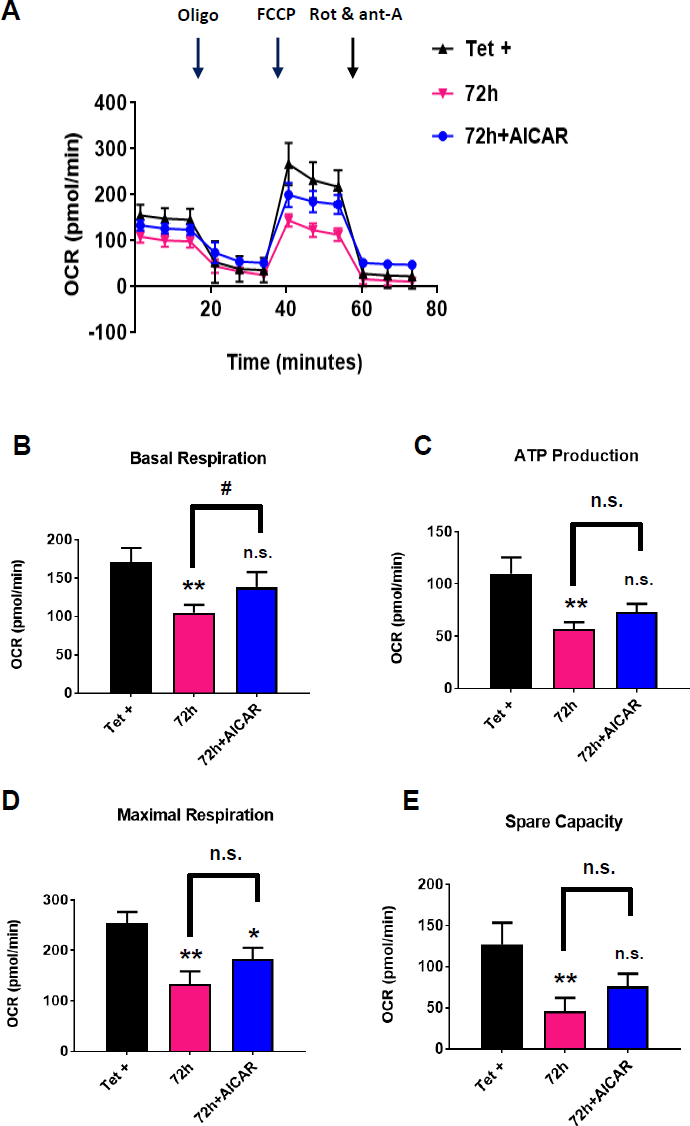
Activation of SIRT3 by AICAR rescues mitochondrial dysfunction induced by αsyn oligomers. (A) Mitochondrial OCR was assessed by Seahorse XFe96 Analyzer. The OCR is significantly reduced in cells overexpressing αsyn and SIRT3 activation significantly improves the OCR levels and respiratory function (n=4). (B) Basal respiration. (C) ATP production (D) maximal respiration (E) spare respiratory capacity. Error bars represent the mean ± S.E.M. *P < 0.05, **P < 0.01 compared to control conditions; #P < 0.05 compared to AICAR treatment. n.s: not significant.

### SIRT3 deficit is also present *in vivo*

Although a very recent study demonstrated that overexpression of SIRT3 in a rodent model of αsyn overexpression could rescue αsyn-induced cell loss in the substantia nigra (SN) pars compacta (Gleave *et al.,* 2017), the mechanism by which SIRT3 exerts its neuroprotective effects was not addressed. To confirm the findings of the previous study and determine if similar mechanisms are at play *in vivo* to those described in our cellular studies, we used a rodent model whereby accumulation of αsyn oligomers in the SN and striatum (STR) after 4 weeks is accompanied by significant loss of dopaminergic neurons. We have previously shown that unilateral injection of AAV2/8-human αsyn into SN of adult rat results in abundant expression of αsyn oligomeric species in both cell bodies and axon terminals of the nigrostriatal pathway (Delenclos *et al.,* 2016). Here, we performed cellular fractionation of nigral and striatal tissue 4 weeks after viral transduction and assessed cytosolic and mitochondrial fractions for SIRT3 levels in the ipsilateral (injected) side of the brain. Consistent with our *in vitro* data, accumulation of αsyn in SN was accompanied by a significant decrease in SIRT3 protein levels (Fig. 6A, **P < 0.01). At this time point no differences were detected in SIRT3 levels between the ipsilateral and control/uninjected side in the STR. Of note, there was no difference in SIRT3 levels in SN in control animals that received an injection of AAV8 expressing gaussia luciferase only (Suppl. Fig. 4A). Furthermore, examination of OPA1 and DRP1 levels in our rodent model revealed results consistent with our in vitro data, with DRP1 significantly increased in the injected SN (Fig. 6B, *P<0.05) and OPA1 significantly decreased (Fig. 6B, *P<0.05). Lastly, AMPKα-CREB signaling was downregulated in the SN of these animals (Suppl. Fig. 4B), mimicking once again our *in vitro* observation.

**Figure 6:**
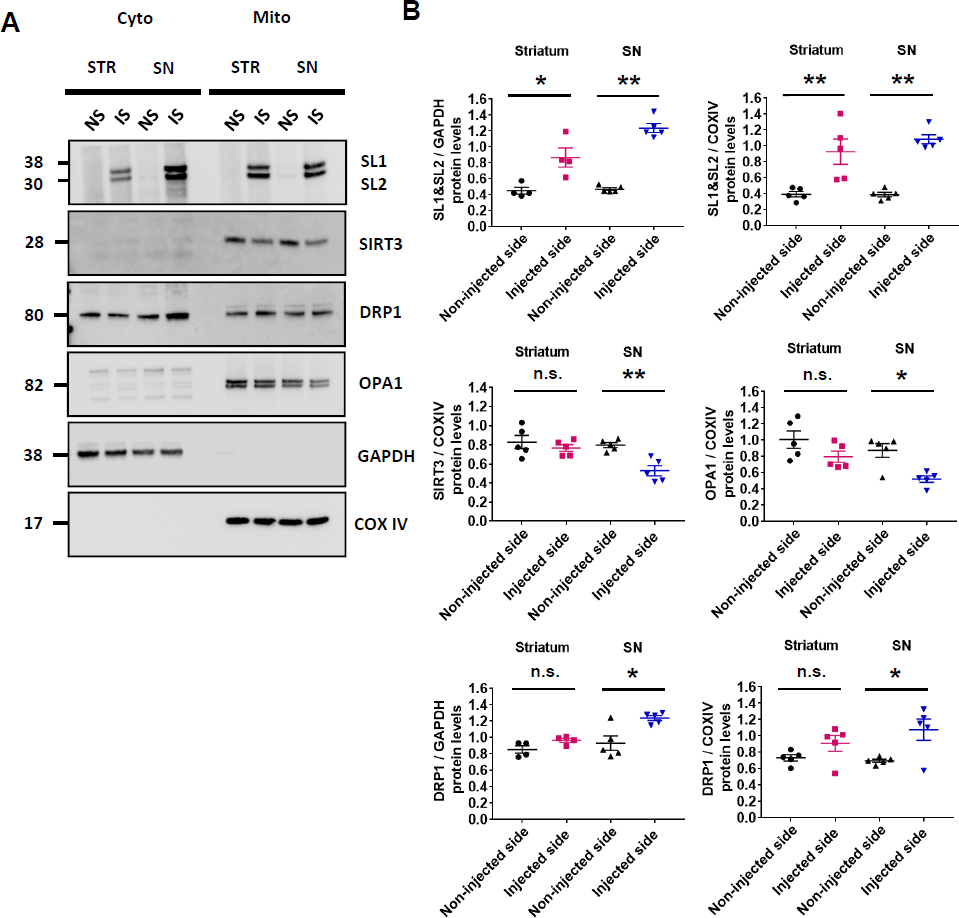
αSyn induces SIRT3 inactivation and modifies normal mitochondrial dynamics *in vivo*. (A) Representative cropped western blots showing αsyn, SIRT3, DRP1, and OPA1 in cytosol and mitochondria from STR and SN of two rats injected with AAV8-SL1 and AAV8-SL2 after 4 weeks. αSyn expression increases DRP1 and decreases SIRT3 and OPA1 levels in injected side (IS) compared to non-injected side (NS). (B) Quantification of αsyn, SIRT3, DRP1, and OPA1 protein levels in cytosol and mitochondria from two separate blots for each of 4-5 rats. All bands were normalized to respective loading controls GAPDH and COXIV. In panel (A) the same samples were run on one blot that was cropped prior to immunoblotting for αsyn, SIRT3, COXIV, DRP1, OPA1, and GAPDH. Error bars represent the mean ± S.E.M (n = 4-5 rats). *P < 0.05, **P < 0.01 compared to control conditions. n.s: not significant.

### SIRT3 levels are decreased in human Lewy body disease brains

Lastly, we assessed the level of SIRT3 in human post mortem brain with a confirmed neuropathological diagnosis of Lewy body disease (LBD) (Table 1). Frozen striatal tissue from 10 LBD and 10 healthy controls was homogenized, run on SDS-PAGE, and probed with antibodies to detect SIRT3, OPA1, and DRP1. Western blot analyses showed significantly reduced expression of SIRT3 in LBD brains compared to controls (Fig. 7A and B). We also detected reduced expression of OPA1 protein but no significant difference in the level of DRP1 compared to controls (Fig. 7A and B). Brains from both sexes were utilized but there was no difference in the interpretation of the data when stratified by sex (data not shown). Together, these results are consistent with our findings from cell and rodent models with decreased SIRT3 protein levels when αsyn accumulates and aggregates in neurons.

**Figure 7:**
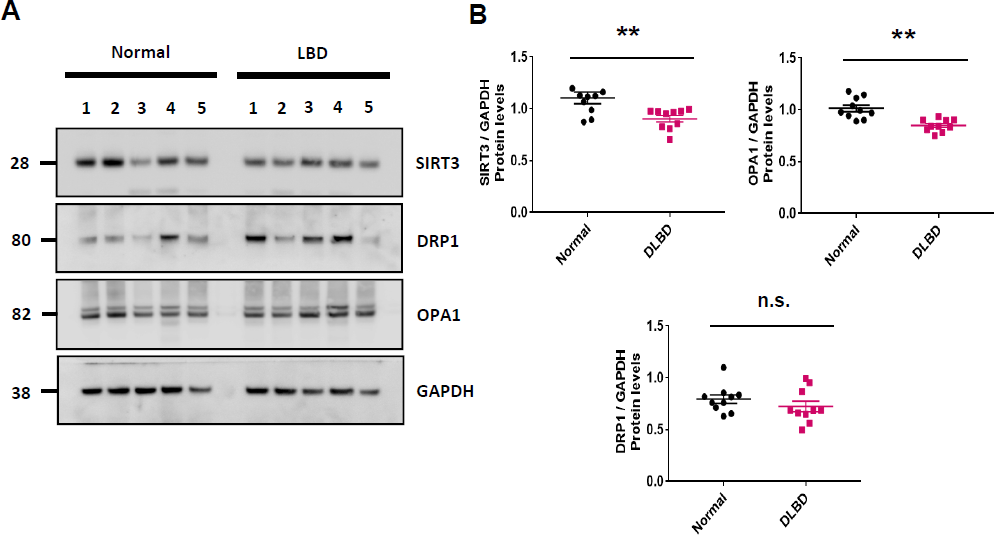
SIRT3 is decreased in human post-mortem brain of neuropathologically confirmed Lewy body disease individuals. (A) Representative cropped western blot from n=3 showing decreased SIRT3, and OPA1 in human post mortem brain of five Lewy body disease (LBD) brains compared to five controls. No significant difference in DRP1 protein levels was detected. (B) Quantification of SIRT3, DRP1, and OPA1 protein levels from n=3 western blots of whole brain lysates from ten LBD and ten control brains normalized to GAPDH loading control. Error bars represent the mean ± S.E.M. **P < 0.01 compared to control conditions. n.s: not significant.

**Figure 8:**
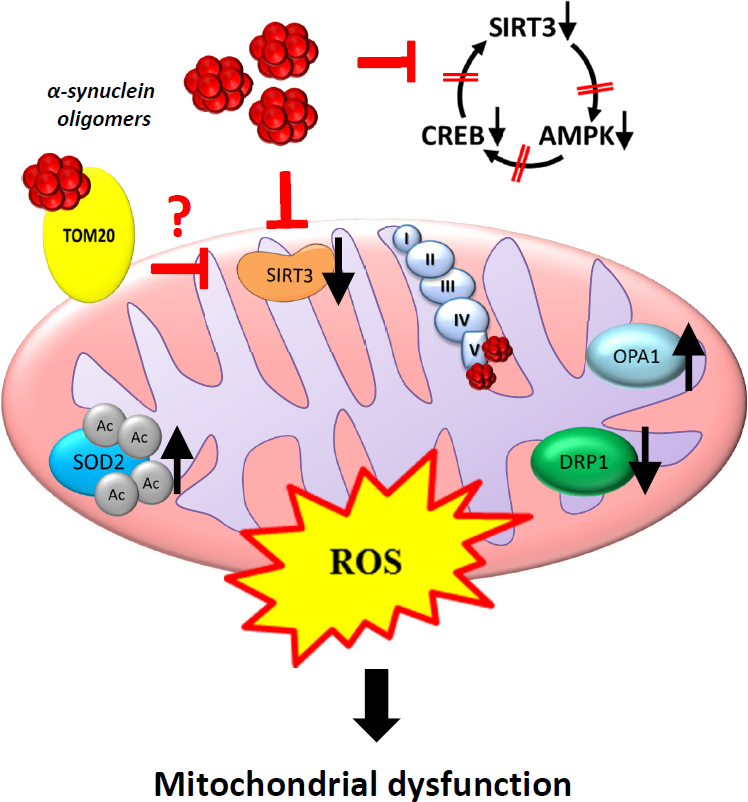
A schematic illustration of potential mechanisms αsyn-induced SIRT3 inactivation and mitochondrial dysfunction. A speculative mechanism whereby mitochondrial αsyn oligomers reduce mitochondrial SIRT3 levels is impaired translocation of SIRT3 from cytosol due to αsyn/TOM20 interaction (Di Maio et al. 2016). Consequences of decreased SIRT3 include decreased AMPK-CREB signaling, impairment in mitochondrial bioenergetics and dynamics, and increased acetylation of SIRT3 substrates such as SOD2 all of which contribute to increased ROS production and neurodegeneration. Question mark indicates pathway not supported by data in this manuscript.

## Discussion

Herein, we identify a cellular mechanism that explains how mitochondrial αsyn oligomers lead to mitochondrial dysfunction and the initiation of a self-perpetuating cycle of aggregation and deficient cellular metabolism that eventually results in cell death. For the first time we identify decreased SIRT3 activity as a consequence of αsyn oligomer accumulation in mitochondria in multiple model systems including cell models, animal models, and human postmortem brains with a neuropathological diagnosis of LBD. We demonstrate the presence of αsyn oligomers in mitochondria correlate with decreased mitochondrial function and decreased SIRT3 expression and function. Interestingly, we show that SIRT3 downregulation is accompanied by dysregulation of AMPK signaling pathway, perturbation of fusion/fission mechanisms, and impairment of basal respiration, all of which contribute to increase ROS and mitochondrial dysfunction. These findings are observed not only in an experimental cellular model, but also in a rodent model of αsyn aggregation, and more importantly, in human post mortem LBD brain. Lastly, treatment with an AMPK agonist, AICAR, improves αsyn-induced mitochondrial dysfunction by restoring SIRT3 expression and decreasing αsyn oligomer formation. Overall, these results demonstrate the health enhancing capabilities of SIRT3 and validate its potential as a new therapeutic target for Parkinson disease and related disorders.

Mitochondrial dysfunction has been linked to the pathogenesis of neurodegenerative diseases including Parkinson disease, with mutations identified in mitochondrial-associated proteins such as PINK1 and parkin causing familial Parkinson disease (Schapira *et al.,* 1993; Dawson *et al.,* 2003). αSyn, a major neuropathological hallmark of Parkinson disease and alpha-synucleinopathies can perturb mitochondria and previous studies have shown that overexpression of αsyn has dramatic effects on mitochondrial morphology, reduces respiratory chain complex activity, and impairs mitochondrial functions *in vitro* and *in vivo* (Siddiqui *et al.,* 2012; Bobela *et al.,* 2017). Accumulation of wild-type αsyn and truncated species within the mitochondria has been described (Sarafian *et al.,* 2013; Subramaniam *et al.,* 2014) however, no study has definitively demonstrated the presence of oligomeric αsyn species in mitochondria. Here, we used a split luciferase protein complementation assay to demonstrate accumulation of αsyn oligomers in the mitochondrial fraction of cells in culture and in rat brain homogenates. We speculate that the presence of oligomeric αsyn species triggers a cascade of events leading to mitochondria malfunction associated with Parkinson disease pathogenesis. Deficiency of SIRT3 is observed in cellular models of Huntington’s disease (Fu *et al.,* 2012) and down regulation of SIRT3 increases dopaminergic cell death in an MPTP mouse model of Parkinson disease (Liu *et al.,* 2015). Most recently, overexpression of SIRT3 was demonstrated to prevent αsyn-induced neurodegeneration in a rodent AAV model (Gleave *et al.,* 2017). In humans, down downregulation of SIRT3 has been previously reported in post-mortem human Alzheimer disease brain (Han *et al.,* 2014; Lee *et al.,* 2018).

SIRT3 is emerging as an important regulator of cellular biogenesis and oxidative stress. Recent evidence supports attenuation of ROS and improved mitochondrial bioenergetics upon activation of SIRT3 (Ramesh *et al.,* 2018), while SIRT3 knockdown exacerbates ROS production (Zhang *et al.,* 2016). The current school of thought is that SIRT3 induces neuroprotection by enhancing mitochondrial biogenesis and integrity, perhaps by increasing mitochondrial DNA content and suppressing SOD activity (Dai *et al.,* 2014a,b; Zhang *et al.,* 2016; Liu *et al.,* 2017). AMPK is upstream of SIRT3 in the signaling pathway that regulates gene expression and the activity of nicotinamide phosphoribosyl transferase (NAMPT) (Fulco *et al.,* 2008; Costford *et al.,* 2010). SIRT3 also seems to be under the control of AMPK/CREB-PGC-1α signaling pathway known to have a crucial role in the regulation of mitochondrial biogenesis and function, activating mitochondrial enzymes involved in antioxidant defenses and metabolism (Shi *et al.,* 2005; Kong *et al.,* 2010; Abdel Khalek *et al.,* 2014). Here, we tested the hypothesis that the AMPK/CREB signaling pathway plays an important role in αsyn-induced SIRT3 down-regulation. Overexpression of oligomeric αsyn significantly decreased levels of p-AMPKα and p-CREB both *in vitro* and *in vivo.* Moreover, pharmacological activation of AMPK by AICAR was able to restore levels of p-AMPKα and p-CREB, and increase mitochondrial SIRT3 protein expression. Most importantly, we found that activating AMPK significantly reduced the level of αsyn oligomers in our cellular model system. These data raise the question of whether modulating SIRT3 levels will alleviate αsyn-induced pathology and slow or halt αsyn-induced cellular dysfunction in Parkinson disease and related synucleinopathies.

Mitochondria are dynamic organelles that continuously undergo fission and fusion, processes necessary for cell survival and adaptation to changing energy requirements for cell growth, division, and distribution of mitochondria during differentiation (van der Bliek *et al.,* 2013). Our results demonstrate that mitochondrial dynamics are modified by the presence of αsyn oligomers in mitochondria. Impaired fission/fusion balance is demonstrated herein by reduced levels of OPA1, and increased DRP1 and phosphorylated DRP1. Under stress conditions, DRP1 is recruited to mitochondria where it initiates mitochondrial fission and induces mitochondrial dysfunction. DRP1 activity is regulated by several post-translational modifications including phosphorylation at Serine 616 (Elgass *et al.,* 2013), which rapidly activates DRP1 and stimulates mitochondrial fission during mitosis (Cho *et al.,* 2013; Sanchis-Gomar and Derbré, 2014). When αsyn oligomers localize to mitochondria, we observe an accompanying decrease in OPA1, driving mitochondria dynamics toward fission and fragmentation, indicating that αsyn oligomers induce mitochondrial dysfunction by regulating mitochondrial dynamics. AICAR-treatment was able to restore OPA1 and DRP1 protein expression to control levels, subsequently resulting in improved mitochondrial function, indicating that SIRT3 plays an important role in regulating of maintenance of mitochondrial function during stress.

Our results identify a mechanism whereby mitochondrial αsyn oligomers contribute to impaired mitochondrial respiration and impaired mitochondrial dynamics by disrupting AMPK/CREB/SIRT3 signaling. These data are consistent with a very recent study demonstrating interaction of αsyn with ATP synthase in the mitochondria and impairment of complex I-dependent respiration (Ludtmann et al, 2018). Additionally, previous studies have shown that αsyn can also interact with TOM20 (Di Maio *et al.,* 2016), which is required for mitochondrial protein import, and decrease its function. SIRT3 is reported to exist in the cytoplasm in an inactive form and recruited to the mitochondria upon stress (Anamika *et al.,* 2017). It is tempting to speculate that TOM20 plays a role in the translocation of SIRT3 to mitochondria and that αsyn-induced deficit in protein import result in reduced mitochondrial SIRT3 levels thereby initiating the cascade of mitochondrial dysfunction that results in decreased mitochondrial bioenergetics. Further studies will be necessary to determine if there is any substance to this speculation and additional studies should address the role of SIRT3 deacetylation substrates as possible players in Parkinson disease pathogenesis. Taken together, our study opens the door to the use of SIRT3 activators as potential therapeutics for restoration of mitochondrial deficits and decrease in αsyn-induced pathophysiology.

## Acknowledgements

We thank Dr. Dennis Dickson, Dr. Michael DeTure, and the Mayo Clinic Brain bank for human post-mortem brain samples used in this study.

## References

Abdel Khalek W, Cortade F, Ollendorff V, Lapasset L, Tintignac L, Chabi B, et al. SIRT3, a mitochondrial NAD+-dependent deacetylase, is involved in the regulation of myoblast differentiation. PLoS One 2014; 9:e114388.

Anamika, Khanna A, Acharjee P, Acharjee A, Trigun SK. Mitochondrial SIRT3 and neurodegenerative brain disorders. [Review]. J Chem Neuroanat 2017; In Press.

Ansari A, Rahman MS, Saha SK, Saikot FK, Deep A, Kim KH. Function of the SIRT3 mitochondrial deacetylase in cellular physiology, cancer, and neurodegenerative disease. Aging Cell 2017; 16: 4–16.

Bansal S, Biswas G, Avadhani NG. Mitochondria-targeted heme oxygenase-1 induces oxidative stress and mitochondrial dysfunction in macrophages, kidney fibroblasts and in chronic alcohol heaptotoxicity. Redox Biol 2013; 2: 273–83.

Bause AS, Haigis MC. SIRT3 regulation of mitochondrial oxidative stress. Exp Gerontol 2013; 48: 634–39.

Bobela W, Nazeeruddin S, Knott G, Aebischer P, Schneider BL. Modulating the catalytic activity of AMPK has neuroprotective effects against α-synuclein toxicity. Mol Neurodegener 2017; 12: 80.

Cho B, Choi SY, Cho HM, Kim HJ, Sun W. Physiological and pathological significance of dynamin- related protein 1 (Drp1)-dependent mitochondrial fission in the nervous system. Exp Neurobiol 2013; 22: 149–57.

Costford SR, Bajpeyi S, Pasarica M, Albarado DC, Thomas SC, Xie H, Church TS, Jubrias SA, Conley KE, Smith SR. Skeletal muscle NAMPT is induced by exercise in humans. Am J Physiol Endocrinol Metab 2010; 298: E117–26.

Dai SH, Chen T, Wang YH, Zhu J, Luo P, Rao W, et al. Sirt3 protects cortical neurons against oxidative stress via regulating mitochondrial Ca2^+^ and mitochondrial biogenesis. Int J Mol Sci 2014a; 15: 14591–609.

Dai SH, Chen T, Wang YH, Zhu J, Luo P, Rao W, et al. Sirt3 attenuates hydrogen peroxide induced oxidative stress through the preservation of mitochondrial function in HT22 cells. Int J Mol Med. 2014b; 34:1159–68.

Dawson TM, Dawson VL. Molecular pathways of neurodegeneration in Parkinson’s disease. [Review]. Science 2003; 302: 819–22.

Delenclos M, Trendafilova T, Jones DR, Moussaud S, Baine AM, Yue M, et al. A Rapid, Semi-Quantitative Assay to Screen for Modulators of Alpha-Synuclein Oligomerization Ex vivo. Front Neurosci 2016; 9: 511.

Devi L, Raghavendran V, Prabhu BM, Avadhani NG, Anandatheerthavarada HK. Mitochondrial import and accumulation of alpha-synuclein impair complex I in human dopaminergic neuronal cultures and Parkinson disease brain. J Biol Chem 2008; 283:9089–100.

Di Maio R, Barrett PJ, Hoffman EK, Barrett CW, Zharikov A, Borah A, et al. α-Synuclein binds to TOM20 and inhibits mitochondrial protein import in Parkinson’s disease. Sci Transl 2016; 8: 342ra78.

Dulovic M, Jovanovic M, Xilouri M, Stefanis L, Harhaji-Trajkovic L, Kravic-Stevovic T, et al. The protective role of AMP-activated protein kinase in alpha-synuclein neurotoxicity in vitro. Neurobiol Dis 2014; 63: 1–11.

Elgass K, Pakay J, Ryan MT, Palmer CS. Recent advances into the understanding of mitochondrial fission. Biochim Biophys Acta 2013; 1833: 150–61.

Fu J, Jin J, Cichewicz RH, Hageman SA, Ellis TK, Xiang L, et al. Trans-(-)-ε-Viniferin increases mitochondrial sirtuin 3 (SIRT3), activates AMP-activated protein kinase (AMPK), and protects cells in models of Huntington Disease. J Biol Chem 2012; 287:24460–72.

Fulco M, Cen Y, Zhao P, Hoffman EP, McBurney MW, Sauve AA, et al. Glucose restriction inhibits skeletal myoblast differentiation by activating SIRT1 through AMPK-mediated regulation of Nampt. Dev Cell 2008; 14: 661–73.

Gleave JA, Arathoon LR, Trinh D, Lizal KE, Giguère N, Barber JHM, et al. Sirtuin 3 rescues neurons through the stabilisation of mitochondrial biogenetics in the virally-expressing mutant α-synuclein rat model of parkinsonism. Neurobiol Dis 2017; 106: 133–46.

Han P, Tang Z, Yin J, Maalouf M, Beach TG, Reiman EM, et al. Pituitary adenylate cyclase-activating polypeptide protects against β-amyloid toxicity. Neurobiol Aging 2014; 35:2064–71.

Hebert AS, Dittenhafer-Reed KE, Yu W, Bailey DJ, Selen ES, Boersma MD, et al. Calorie restriction and SIRT3 trigger global reprogramming of the mitochondrial protein acety-lome. Mol Cell 2013; 49: 186–99.

Herskovits AZ, Guarente L. Sirtuin deacetylases in neurodegenerative diseases of aging. [Review]. Cell Res 2013; 23: 746–58.

Hsu LJ, Sagara Y, Arroyo A, Rockenstein E, Sisk A, Mallory M, et al. alpha-synuclein promotes mitochondrial deficit and oxidative stress. Am J Pathol 2000; 157: 401–10.

Kim SH, Lu HF, Alano CC. Neuronal Sirt3 protects against excitotoxic injury in mouse cortical neuron culture. PLoS One 2011; 6: e14731.

Kong X, Wang R, Xue Y, Liu X, Zhang H, Chen Y, et al. Sirtuin 3 a new target of PGC-1α, plays an important role in the suppression of ROS and mitochondrial biogenesis. PLoS One 2010; 5: e11707.

Kyrylenko S, Baniahmad A. Sirtuin family: a link to metabolic signaling and senescence. Curr Med Chem 2010; 17: 2921–32.

Lee J, Kim Y, Liu T, Hwang YJ, Hyeon SJ, Im H, et al. SIRT3 deregulation is linked to mitochondrial dysfunction in Alzheimer’s disease. Aging Cell 2018; 17: doi: 10.1111/acel.12679.

Liu J, Li D, Zhang T, Tong Q, Ye RD, Lin L. SIRT3 protects hepatocytes from oxidative injury by enhancing ROS scavenging and mitochondrial integrity. Cell Death Dis 2017; 8:e3158.

Liu L, Peritore C, Ginsberg J, Kayhan M, Donmez G. SIRT3 attenuates MPTP-induced nigrostriatal degeneration via enhancing mitochondrial antioxidant capacity. Neurochem Res 2015; 40: 600–8.

Lombard DB, Alt FW, Cheng HL, Bunkenborg J, Streeper RS, Mostoslavsky R, et al. Mammalian Sir2 homolog SIRT3 regulates global mitochondrial lysine acetylation. Mol Cell Biol 2007; 27: 8807–14.

López-Otín C, Blasco MA, Partridge L, Serrano M, Kroemer G. The hallmarks of aging. [Review]. Cell 2013; 153: 1194–217.

Ludtmann MHR, Angelova PR, Horrocks MH, Choi ML, Rodrigues M, Baev AY, et al. α-Synuclein oligomers interact with ATP synthase and open the permeability transition pore in Parkinson’s disease. Nat Commun. 2018 Jun 12;9(1):2293. doi: 10.1038/s41467-018-04422-2.

Marongiu R, Spencer B, Crews L, Adame A, Patrick C, Trejo M, et al. Mutant Pink1 induces mitochondrial dysfunction in a neuronal cell model of Parkinson’s disease by disturbing calcium flux. J Neurochem 2009; 108: 1561–74.

Moussaud S, Malany S, Mehta A, Vasile S, Smith LH, McLean PJ. Targeting α-synuclein oligomers by protein-fragment complementation for drug discovery in synucleinopathies. Expert Opin Ther Targets 2015; 19: 589–603.

Nakamura K, Nemani VM, Azarbal F, Skibinski G, Levy JM, Egami K, et al. Direct membrane association drives mitochondrial fission by the Parkinson disease-associated protein alpha-synuclein. J Biol Chem 2011; 286: 20710–26.

Paxinos G, Watson C. The Rat Brain in Stereotaxic Coordinates (4th ed). San Diego, CA: Academic Press; 1998.

Pillai VB, Sundaresan NR, Kim G, Gupta M, Rajamohan SB, Pillai JB, et al. Exogenous NAD blocks cardiac hypertrophic response via activation of the SIRT3-LKB1-AMP-activated kinase pathway. J Biol Chem 2010; 285: 133–3144.

Qiu X, Brown K, Hirschey MD, Verdin E, Chen D. Calorie restriction reduces oxidative stress by SIRT3-mediated SOD2 activation. Cell Metab 2010; 12: 662–7.

Ramesh S, Govindarajulu M, Lynd T, Briggs G, Adamek D, Jones E, et al. SIRT3 activator Honokiol attenuates β-Amyloid by modulating amyloidogenic pathway. PLoS One 2018;13: e0190350.

Reeve AK, Ludtmann MH, Angelova PR, Simcox EM, Horrocks MH, Klenerman D, et al. Aggregated α-synuclein and complex I deficiency: exploration of their relationship in differentiated neurons. Cell Death Dis 2015; 6: e1820.

Sanchis-Gomar F, Derbré F. Mitochondrial fission and fusion in human diseases. N Engl J Med 2014; 370: 1073–74.

Sarafian TA, Ryan CM, Souda P, Masliah E, Kar UK, Vinters HV, et al. Impairment of mitochondria in adult mouse brain overexpressing predominantly full-length, N-terminally acetylated human α-synuclein. PLoS One 2013; 8: e63557.

Shipper HM, Liberman A, Stopa EG. Neural heme oxygenase-1 expression in idiopathic Parkinson’s disease. Exp Neurol 1998; 150: 60–8.

Schapira AH, Hartley A, Cleeter MW, Cooper JM. Free radicals and mitochondrial dysfunction in Parkinson’s disease. [Review]. Biochem Soc Trans 1993; 21: 367–70.

Shi T, Wang F, Stieren E, Tong Q. SIRT3, a mitochondrial sirtuin deacetylase, regulates mitochondrial function and thermogenesis in brown adipocytes. J Biol Chem 2005; 280:13560–67.

Siddiqui A, Chinta SJ, Mallajosyula JK, Rajagopolan S, Hanson I, Rane A, et al. Selective binding of nuclear alpha-synuclein to the PGC1alpha promoter under conditions of oxidative stress may contribute to losses in mitochondrial function: implications for Parkinson’s disease. Free Radic Biol Med 2012; 53: 993–1003.

Song W, Patel A, Qureshi HY, Han D, Schipper HM, Paudel HK. The Parkinson disease-associated A30P mutation stabilizes alpha-synuclein against proteasomal degradation triggered by heme oxygenase-1 over-expression in human neuroblastoma cells. J Neurochem 2009; 110: 719–33.

Subramaniam SR, Vergnes L, Franich NR, Reue K, Chesselet MF. Region specific mitochondrial impairment in mice with widespread overexpression of alpha-synuclein. Neurobiol Dis 2014; 70: 204–13.

Takeuchi K, Morizane Y, Kamami-Levy C, Suzuki J, Kayama M, Cai W, et al. AMP-dependent kinase inhibits oxidative stress-induced caveolin-1 phosphorylation and endocytosis by suppressing the dissociation between c-Abl and Prdx1 proteins in endothelial cells. J Biol Chem 2013; 288: 20581–591.

Torrens-Mas M, Oliver J, Roca P, Sastre-Serra J. SIRT3: Oncogene and Tumor Suppressor in Cancer. [Review]. Cancers (Basel) 2017; 9: 90.

van der Bliek AM, Shen Q, Kawajiri S. Mechanisms of mitochondrial fission and fusion. [Review]. Cold Spring Harb Perspect Biol 2013; 5: a011072.

Weir HJ, Murray TK, Kehoe PG, Love S, Verdin EM, O’Neill MJ, et al. CNS SIRT3 expression is altered by reactive oxygen species and in Alzheimer’s disease. PLoS One 2012; 7: e48225.

Yin J, Han P, Tang Z, Liu Q, Shi J. Sirtuin 3 mediates neuroprotection of ketones against ischemic stroke. J Cereb Blood Flow Metab 2015; 35: 1783–89.

Zhang JY, Deng YN, Zhang M, Su H, Qu QM. SIRT3 Acts as a Neuroprotective Agent in Rotenone-Induced Parkinson Cell Model. Neurochem Res 2016; 41: 1761–73.

